# Single-Cell proteomics discerns patient-specific subpopulations in pediatric B-cell acute lymphoblastic leukemia

**DOI:** 10.64898/2026.08.27.747627

**Authors:** Farah Jayousi, Felix Kraus, Agustina Conrrero, Jonathan W Bush, Audi Setiadi, Philipp F Lange

## Abstract

B-cell acute lymphoblastic leukemia (B-ALL) is the most common childhood cancer, representing ∼30% of pediatric cancers and ∼80-85% of pediatric ALL cases. Despite high remission rates, relapse remains a major challenge, often driven by therapy-resistant subpopulations, which are masked in bulk analysis, thus limiting our understanding of disease progression and optimal therapeutic intervention. Precision oncology enables proteome-level characterization of patient specific cancer samples and their subpopulations, essential for improving prognostic accuracy and individualized therapies. Single-cell proteomics by mass spectrometry (SCP-MS) enables quantification of hundreds to thousands of proteins at single cell level, uncovering cellular programs that may contribute to minimal residual disease (MRD) and relapse. Here, employing single-cell sorting of leukemic cells coupled with high-sensitivity SCP-MS, we profile individual leukemic blasts and normal immature B-cells from pediatric B-ALL bone marrow aspirates alongside age-matched controls. SCP-MS deconvoluted cellular heterogeneity and revealed subpopulations with variable leukemia-marker expression, highlighting its potential for early detection of relapse-prone phenotypes and personalized pediatric therapy.

## Introduction

Acute lymphoblastic leukemia (ALL) is a malignancy of lymphoid progenitor cells, where the aberrant proliferation of immature lymphoblasts results in the disruption of ‘normal’ hematopoiesis. ALL is classified by lineage (B-cell ALL; pediatric cases: 85%, adult cases: 75% / T-cell ALL; pediatric cases: 15%, adult cases 25%) and stratification into pediatric and adult is based on average age of peak incidence rates (under five years for pediatric and 53 years for adult)^1^. B-ALL is the most common childhood malignancy, accounting for ∼30% of pediatric cancers^4^and ∼80–85% of pediatric ALL cases^5^. Despite high remission rates with frontline therapies, relapse remains a leading cause of mortality in children with B-ALL, often driven by the emergence of unique subpopulations originating from rare, therapy-resistant leukemic cells^6^.

There are significant and notable differences between adult and pediatric relapses in ALL, mainly the more stable, and thus more predictable, genetic rearrangements (e.g. ETV6:RUNX1) present in pediatric cases^2,3^. Despite the stable genetic re-arrangements, predictions of relapse are hard; clinicians and researchers require a deeper understanding of the biological factors contributing to either full remission or clinical relapse^2,3^.

A key challenging factor influencing prognosis and outcome in children is the fragmentation of the B-ALL molecular Jayousi et al. 2026 (BioRvix-preprint) subtypes and intra-patient heterogeneity. There are over 20 classified molecular profiles of pediatric B-ALL and this disparity affects treatment responsiveness and long-term survival^7^. For example, hyperdiploidy is the most common subtype in pediatric B-ALL and is associated with excellent prognosis^8^. However, a genetic fusion such as the ***BCR-ABL1*** subtype is linked with a higher likelihood of relapse, poor prognosis and therapy resistance^7,9^. The molecular etiology responsible for this disease heterogeneity across pediatric B-ALL patients remains elusive^7–9^.

Currently, the bulk of clinical characterization relies on a panel of complementary techniques, including cytogenetics for characterization of chromosomal re-arrangements, whole genome and bulk RNA sequencing, pathological assessment, and robust multi-parameter clinical flow-cytometry for the analysis of cellular immunophenotypic features at single cell resolution, albeit lacking the depth of genomic or proteomic approaches.

Over the past decade, it has become increasingly evident that discovering insights to B-ALL relapse prevention requires a deeper understanding of the cellular and molecular heterogeneity and plasticity within the leukemic niche^10,11^. Branched evolution of genetic subclones is well documented^12^and single-cell RNA sequencing studies provide evidence of pre- and post-treatment diversity as well as plasticity and convergence towards less differentiated and more treatment-resistant states, only partially associated with genetic alterations^13–16^. While these studies clearly demonstrate intra-patient heterogeneity, they provide limited functional insight, as numerous studies have shown limited correlation between mRNA and protein abundance at the bulk and single cell level^17–19^. Past studies of protein levels at single-cell resolution in leukemia have employed mass-cytometry or flow-cytometry approaches that are highly informative but limited to measuring tens of predefined proteins^20^. The largest current experimental gap in understanding pediatric B-ALL heterogeneity and treatment resistance is global, untargeted quantification of hundreds or thousands of proteins in single primary patient cells.

Compared to well established methods like genomics and transcriptomics, SCP-MS is a relatively recent development and lags behind in both applications and adoption in basic research and clinical settings. However, it is one of the most rapidly evolving areas of research. Several groups have enhanced and tested new technologies to advance the field, such as new cleanup methods, optimization of MS parameters for higher throughput, and most importantly, new instruments with ultra-high sensitivity for maximal detection at the small scale^21^. These advances in SCP-MS have now enabled the characterization of proteomic signatures within individual cells and the definition of new cell populations in different biological and disease contexts^22^. For example, SCP-MS has been used to resolve the cellular hierarchy of primary human CD34^+^hematopoietic stem and progenitor cells (HSPCs), identifying stage-specific proteins governing HSPC function and differentiation that were not predicted by their corresponding mRNA transcripts^23^. SCP-MS approaches are increasingly used to resolve intra-tumoral heterogeneity, immune cell states, and drug-resistant subpopulations in solid tumors that are not distinguishable by bulk or transcriptomic profiling alone^25^. In leukemia specifically, SCP-MS was first applied to characterize the cellular hierarchy of an Acute Myeloid Leukemia (AML) culture model, distinguishing leukemic stem cells, progenitors, and blasts by global (untargeted) proteome profiling rather than a pre-defined antibody panel^24^. However, to date B-ALL has not benefited from an investigation by SCP-MS as the particularly small size and low protein content of B-cells^25^and B-ALL leukemic blasts which is approximately 1/8^th^of an epithelial cell^26,27^and 1/6^th^of AML blasts^28^ makes its analysis particularly challenging.

B-ALL is shaped by extensive genetic, transcriptional, and cellular heterogeneity, yet the functional protein-level correlate of this heterogeneity remains largely uncharacterized. Existing single-cell approaches in leukemia have either been restricted to a small, pre-defined panel of proteins or have relied on transcript abundance as an incomplete proxy for the functional state of a cell. At the same time, recent technological advances have established SCP-MS as a feasible and increasingly powerful tool for resolving cellular hierarchies in both healthy hematopoiesis and malignancy, yet it has not been applied to pediatric B-ALL. In this proof of concept, we address this gap by employing SCP-MS for the discovery of cellular and molecular heterogeneity within and across a cohort of pediatric B-ALL patients. We hypothesize that single-cell proteomics has the potential to uncover clinically relevant and molecularly distinct subpopulations in pediatric B-ALL and other leukemias, providing functional insight beyond what genomic and transcriptomic approaches alone can offer.

## Results

### Robust global profiling of isolated human B-lymphocytes by SCP-MS

To evaluate if SCP-MS is sensitive enough to quantify disease relevant proteins at scale in leukemic blasts isolated from blood and bone marrow (BM) biopsies of children with B-ALL and provides new biological insights, we studied a pilot cohort of three B-ALL patients (age 3, 9, 7, sex: all female, called from here on B-ALL-1,2,3) and age- and sex-matched control participants who did not have cancer. Additionally, bone marrow aspirates were analyzed using established clinical flow-cytometry and bulk proteomic approaches, and bone marrow biopsies were analyzed using immunohistochemistry (IHC) panels to gain spatial information **(Fig. 1A)**. Clinical flow-cytometry analysis of the cohort at diagnosis showed the expected presence of 94%, 89% and 87% CD45^+^leukemic blasts, respectively. The blasts exhibited low side scatter, dim CD45, expressed CD19 and CD10, with dim variable CD20 and heterogeneous CD38 expression. Notably, B-ALL-3 contained a discrete CD38-bright subpopulation comprising approximately 2.4% of total singlet events, further demonstrating phenotypic heterogeneity within the leukemic compartment **(Fig. 1B)**. While traditional bulk proteomics approaches on CD19^+^-sorted patient bone marrow mononuclear cells (BMMCs) did readily differentiate proteome profile differences between patients and identify proteins typically elevated in B-ALL despite, sample heterogeneity within patients including the differentiation of the remaining 6-13% normal immature B-cells was masked by averaging protein intensities across all cells intrinsic to the methodology **(Fig. 1C, Supplementary Data 1)**.

**Figure 1:**
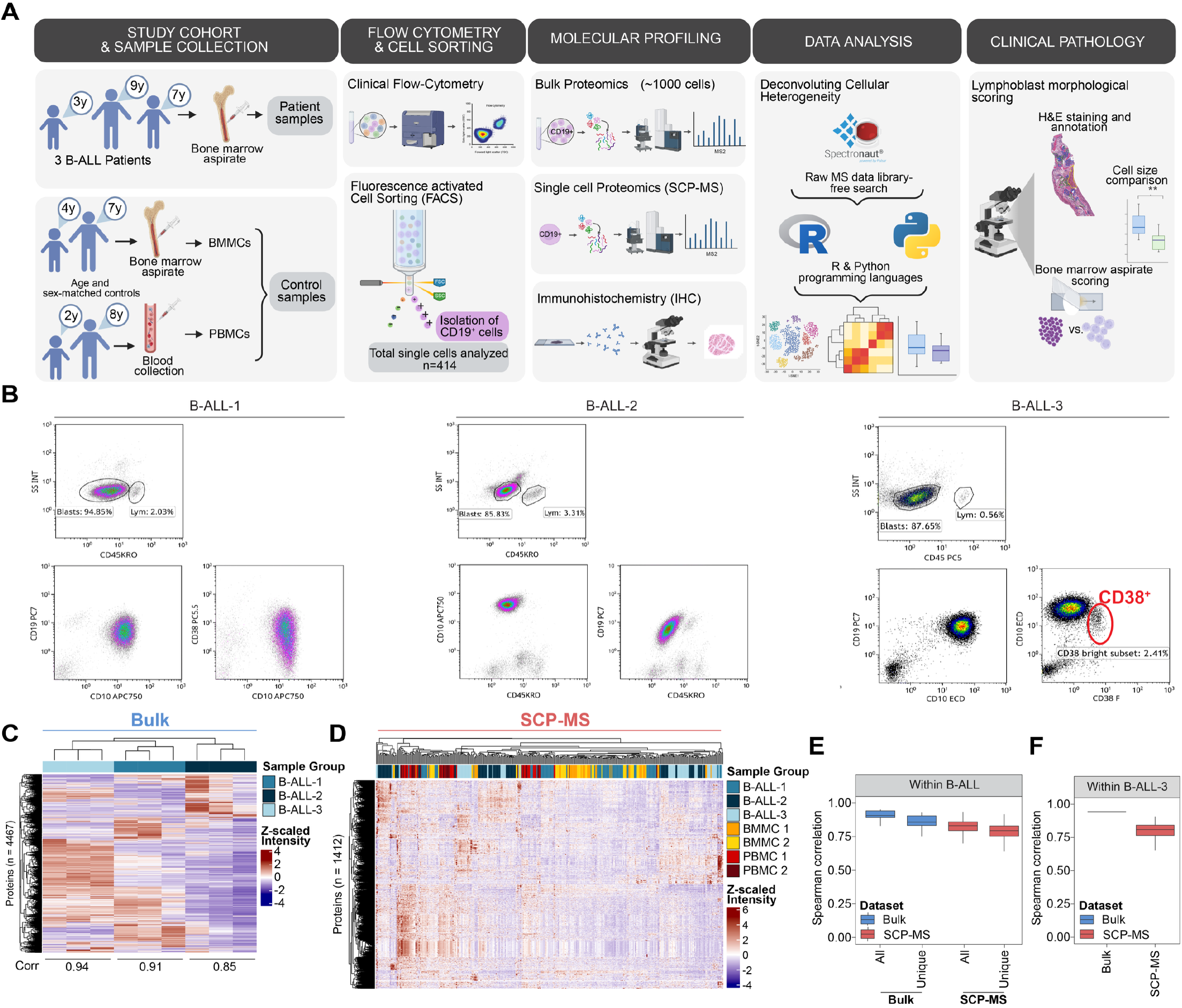
Study overview and initial profiling of pediatric B-ALL patients. **(A)** Schematic overview of the experimental workflow, including patient sample collection, flow-cytometry and fluorescence-activated cell sorting (FACS) for CD19^+^ cell isolation, bulk proteomics, single-cell proteomics (SCP-MS), immunohistochemistry (IHC), and downstream data analysis. **(B)** Clinical flow-cytometry profiles of the three B-ALL patients showing the indicated diagnostic markers and gating strategy for identification and isolation of CD19+ cells. **(C)** Hierarchical clustering heatmap of the three B-ALL patients based on 4,467 proteins, shown as z-scaled protein intensities. **(D)** Hierarchical clustering heatmap of single-cell proteomic profiles based on 1,412 proteins, shown as z-scaled protein intensities. **(E)** Spearman correlation analysis of samples/cells within the same patient, comparing bulk and SCP datasets using all quantified proteins and subsets restricted to proteins quantified exclusively by each method in each patient. **(F)** As E, but for B-ALL-3 alone.

We hypothesized that SCP-MS has the ability to identify any patient-specific subpopulations in an untargeted manner and capture their biological differences at a more granular level than clinical flow cytometry, given its constraint to a limited set of surface proteins. A challenge for SCP-MS in B-ALL patients represent the fact that B-cells have an average median diameter of 6-9µm^29^ and protein content of ∼30-50pg compared to 17-20µm and 150-250pg in HeLa cells a commonly used model for epithelial cells^26,30^.

We used cell lysate dilutions and microfluidic single-cell sorting on HeLa and cultured NALM6 B-cells for consistent ultra-low sample-input while optimizing our SCP-MS method. We modified our MS acquisition method using pydiAID^31^ leading to an increase in proteome depth by 45% (855 vs 1573 protein IDs) with comparable coefficients of variation (%CV) on single-cell equivalent HeLa lysate dilutions **(Supplementary Fig. 1A, Supplementary Data 2**). Next, we injected either 10 single-cell lysates of either HeLa or NCLM6 B-cells in consecutive order and recorded an increase in protein group IDs from a median of 1,954 to 2,644 (∼26% increase in IDs) in normally sized HeLa cells and 994 to 1,542 proteins in cultured NALM6 B-cells (∼35% increase in IDs, **Supplementary Fig. 1B**, Methods). Both methods showed comparable %CV across all quantified proteins in the ten single-cell acquisition runs, supporting robust technical reproducibility (**Supplementary Fig. 1C**).

To increase purity, pediatric cohort samples were stained with FITC-CD19 to separate CD19^+^ B-cells from other peripheral blood mononuclear cells (PBMCs) and other cells in the aspirate using microfluidic sorting prior to SCP-MS analysis. We achieved high isolation accuracy with >90% of wells containing single cells (**Supplementary Fig. 1D**). Utilizing our high-sensitivity pydiAID-optimized acquisition method, we analyzed a total of 414 patient and control participant cells over ∼220 hours of MS analysis time. To monitor instrument stability and performance over the multi-day MS-run time, we injected 250 pg of QC standards (K562 lysates) at the beginning and end of acquisition days (9-day period). Signal intensity was stable over the acquisition period, and we quantified 2855 protein IDs (median; S.D: 48.9), displaying high instrument stability and quantification reliability over time **(Supplementary Fig. 1E)**. We quantified a total of 1,412 proteins across 385 cells passing quality control with a median of 982 proteins per cell and comparable intensity distributions per sample group across the MS-acquisition (**Supplementary Fig. 1F,G**; **Supplementary Data 3**; see Methods). Our SCP-MS workflow showed robust quantitative performance with median %CVs ranging from 38-51% across individual single cells from participants, reflecting biological cell-to-cell variability. Moreover, we assessed cross-sample completeness by calculating the % of protein identification across samples with an average of ∼70% in our single-cell dataset **(Supplementary Fig. 1H,I)**. We evaluated the effect of the experimental setup by mapping the sample group identity and loading order onto a UMAP representation of the SCP-MS data. Embedding of 1,412 detected proteins did resolve sample groups, with a clear separation of B-ALL-1, -2 and -3 cells from both BMMC and PBMC control populations and did not find any clear trajectories influenced by the loading order, suggesting the separation of samples in latent space are driven by the underlying biology and not by technical confounders (**Supplementary Fig. 1J,K**). Quantified peptides across all sample groups did carry high signal intensities confirming that differences in protein IDs between samples were not due to technical detection limits but sample related. **(Supplementary Fig. 1L)**. Together, this demonstrates the feasibility of applying SCP-MS on B-ALL patient samples to obtain high-quality quantitative data across >1,000 proteins to supplement clinical flow-cytometry and enabling deeper molecular profiling for precision oncology.

### Single-cell proteomics identifies heterogenous subpopulations in B-ALL patient samples

Driven by the observation that the analysis of clinical samples using SCP-MS captures a significant fraction of the biological diversity between patients, we aimed to investigate the underlying heterogeneity in more detail. While this SCP-MS workflow demonstrates excellent sensitivity in the context of the limited sample input, it has to be acknowledged that only the top 10% of the theoretical detectable proteome is currently captured. To evaluate if these 10% subset is sufficient to encode key differences between control and malignant cells, as established by bulk studies, we performed differential abundance analysis of all bone marrow-derived patient vs control cells **(Fig. 2A)**. Here we find 29 proteins more abundant and 27 proteins less abundant in B-ALL patients compared with controls, while 1,356 proteins showed no significant changes. Differentially abundant proteins exhibited high median cross-sample completeness coverage (85% and 84% for proteins with decreased and increased abundance, respectively), supporting the robustness of these quantitative differences. Among the differentially abundant proteins, we identified leukemia-associated proteins, including ANP32B, HMGB1 and RAC2.

**Figure 2:**
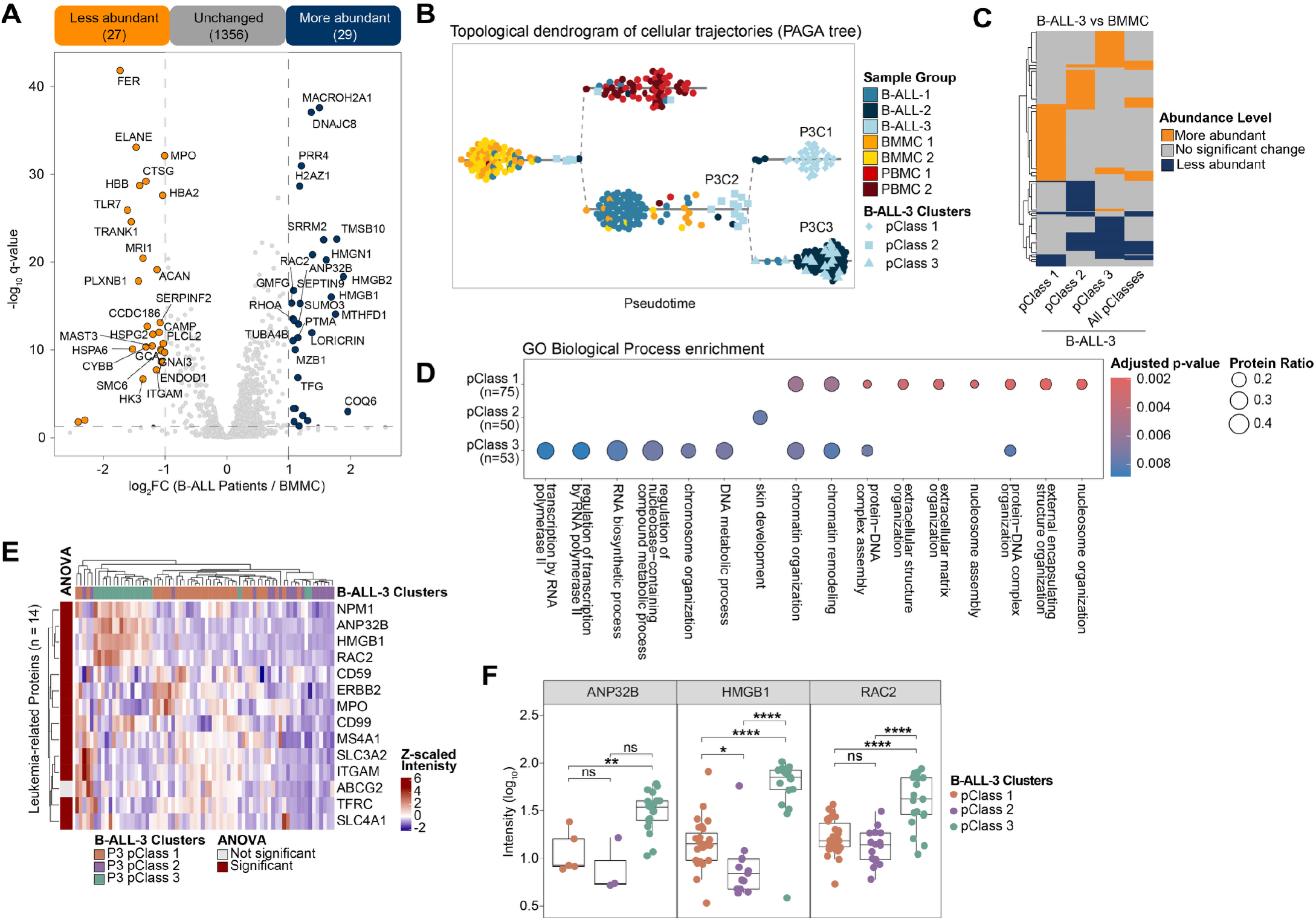
Single-cell proteomic profiling of B-ALL patients and control donors. **(A)** Volcano plot showing differential protein abundance between B-ALL patient cells and BMMC controls (|log_2_FC| > 1, q < 0.05). All significant proteins are highlighted. **(B)** PAGA topological dendrogram depicting inferred cellular trajectories across all conditions. K-means clusters (pClasses) identified within B-ALL-3 are indicated by different shapes. **(C)** Differential protein abundance differences for each B-ALL-3 cluster and all clusters combined relative to BMMC controls. **(D)** Gene Ontology Biological Process enrichment analysis of proteins classified as more abundant in each Patient 3 cluster. Enriched terms are shown according to the adjusted p-value and protein ratio. **(E)** Heatmap of all leukemia-related proteins across B-ALL-3 clusters, with protein intensities shown as z-scaled values. Column colors indicate pClass identity, while row colors indicate ANOVA significance. **(F)** Boxplots of selected, leukemia-related proteins across the three identified pClasses in B-ALL-3.

PAGA (Partition-based Graph Abstraction)^32^trajectory analysis further resolved the relationships between cell populations, placing control BMMCs and PBMCs on distinct branches and identifying three proteomic subpopulations within B-ALL-3 (called pClass 1-3) that were not present in 5 the other patients **(Fig. 2B)** while only one subpopulation was apparent by clinical flow **(Fig 1B)**. Given the unique substructure observed in B-ALL-3, we next characterized the three identified clusters in more detail. Pairwise differential abundance analysis comparing each cluster individually against BMMCs revealed pClass-specific protein signatures that were obscured when all BALL-3 cells were analyzed together, underscoring the importance of resolving intra-patient heterogeneity **(Fig. 2C; Supplementary Fig. 2A-D)**. GO enrichment of the significant protein hits per pClass did reveal unique pathway signatures, especially for pClass-1 and -3, which contained differential signatures of nucleosome and chromatin organization **(Fig. 2D)**. Analysis of selected cancer-associated and leukemia-associated proteins further illustrated differences in leukemic properties across the three clusters **(Fig. 2E,F; Supplementary Fig. 2E)**. This data demonstrates that SCP-MS is able to identify patient-specific subpopulations in an unbiased and robust manner.

### SCP informed clusters map onto distinct spatial organizations in patient biopsies

Among the proteins that most clearly delineated pClass-1 and –3 in B-ALL-3, Nucleolin (NCL) and Mucin-like protein 1 (MUCL1) emerged as leading candidates, with elevated levels particularly in pClass 1 and 3, respectively **(Fig. 3A)**, motivating their selection for orthogonal validation.

**Figure 3:**
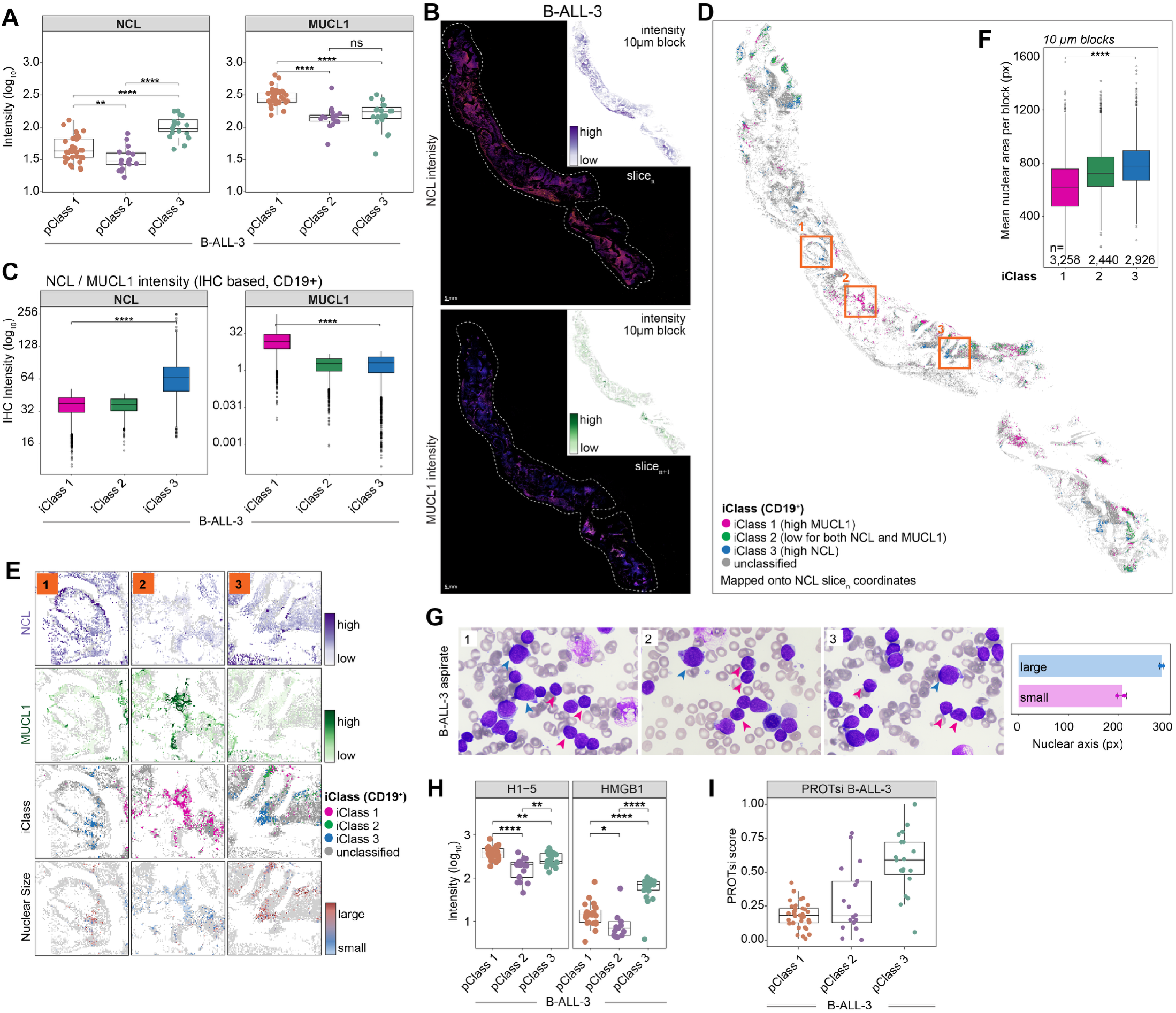
Cross validation of SCP-MS using immunohistochemistry and pathology (previous page). **(A)** Boxplots of NCL (left) and MUCL1 (right) raw proteomics intensity (log_10_) across identified clusters in B-ALL-3 (t-test: *p < 0.05, **p < 0.01, ***p < 0.001, ****p < 0.0001). **(B)** Collage of stitched images of BM section from B-ALL-3 fluorescently labeled against NCL (top) and MUCL1 (bottom). Insets show spatial intensity distribution on the 10 µm block scale. Scale bar: 5 mm. **(C)** Boxplots for NCL (left) and MUCL1 (right) IHC intensity (log_10_) of iClass 1-3 per block. Classes were modeled after SCP-MS abundance of the same proteins (two-sided Wilcoxon test: ****p<0.001). **(D)** Spatial representation of iClass 1-3 distribution onto the BM canvas map. Classes 1-3 are labeled magenta, green and blue. Unclassified blocks are labeled in grey. Orange boxes show the location of zoom-ins from E. **(E)** Zoom-ins marked in D depicting NCL intensity, MUCL1 intensity, iClass assignment and average nuclear size per block (from top to bottom). **(F)** Boxplot of average nuclear area (based on DAPI signal) per cell per class. Number of 10 µm evaluation blocks used for the analysis is shown at the bottom of the plot (****p<0.0001). **(G)** Example microscopy images of aspirate sample from B-ALL-3 and measurement of nuclear axis length (n(large) = 12; n(small) = 28), data from n=3 images. **(H)** Boxplots of select chromatin remodelers H1-5 (left) and HMGB1 (right) raw proteomics intensity (log_10_) across identified clusters in B-ALL-3. **(I)** Boxplot of protein-expression-based stemness index (PROTsi) score for each class in B-ALL-3.

We next performed IHC on BM biopsies of B-ALL-3 against NCL and MUCL1 (consecutive slice_n+1_) to see if we could harmonize and validate our SCP-MS results with IHC data. We co-stained with DAPI and CD19, allowing for nuclear segmentation and cellular classification **(Fig. 3B)**. Using cellposeSAM^33^and custom python scripts, we segmented ∼566k single nuclei together with their spatial information, CD19 status and NCL or MUCL intensity (**Supplementary Fig. 3A**, Methods). Aiming to compare NCL and MUCL1 distributions across cellular morphology, we performed rigid-body and TPS (Thin Plate Spline) alignment to the NCL slice (slice_n_) (**Supplementary Fig. 3B**). Accounting for the uncertainty originating from the spatial shifts between consecutive slices, we rasterized the image space into 10µm blocks for downstream analysis (**Supplementary Fig. 3C**). Modeling the pClass stratification observed by SCP-MS in BALL-3 in our IHC data as iClass-1, -2, -3 (**Fig. 3C**) and mapping these onto the biopsy section (**Fig. 3D**; **Supplementary Fig. 3D**) allowed for the spatial interrogation of proteome signatures in the tissue context. iClasses 1 and 3 did form discrete clusters in distinct regions of the section and showed an apparent correlation with differences in nuclear size (**Fig. 3E**, zoom-ins from orange boxes in **Fig. 3D**, quantified in **Fig. 3F; Supplementary Fig. 3E**). This matched enrichment of nuclear organization-related processes in pClass-1 and -3 as seen by SCP-MS **(Fig. 2D)** and led us to look at the cellular and nuclear morphology in more detail. Morphological assessment by a trained pathologist performed on a representative bone marrow biopsy of B-ALL-3 showed two morphologically distinct blast populations; Blast (1) that are larger in size with finer chromatin and Blast (2) that are smaller in size, with more condensed chromatin **(Supplementary Fig. 3F,G)**. To replicate these results in the liquid biopsy bone marrow aspirate context where the SCP-MS was originally sampled from, we scored aspirate samples for their nuclear morphology **(Fig. 3G)**. In line with our observation on bone marrow biopsies, two distinct populations were observable, with an average nuclear diameter of ∼206 px for small and ∼284 px for large nuclei. Motivated by the nuclear morphology differences observed in IHC, H&E and aspirate samples as well as the GO BP signatures in pClass-1 and -3, we dissected our SCP-MS data in more detail and found drivers of DNA and chromatin condensation differentially expressed between pClass-1 and -3 **(Fig. 3H)**.

Applying protein-expression-based stemness index (PROTsi^34^) on the three subclusters of B-ALL-3, we tested for differences in stemness between these **(Fig. 3I)**. In line with histological and IHC data, pClass-3 did show the highest PROTsi stemness score, suggesting that the observed morphological changes in nuclear size and proteome signatures are linked and describe two different leukemic states. Taken together, these results show that SCP-MS can identify functionally distinct sub-populations that can be cross-validated spatially to further characterize their morphological properties.

## Discussion

In this study, we applied SCP-MS to profile CD19^+^leukemic blasts from pediatric B-ALL patients at the single-cell level. To our knowledge, this represents the first SCP-MS profiling of leukemic blasts directly from pediatric patient samples. Our results demonstrate that SCP-MS can readily resolve proteomic subpopulations within and across patients that are partially masked by patient-driven sample heterogeneity to clinical flow-cytometry and bulk proteomics. Using SCP-MS profiling, we observed that B-ALL-3 of our cohort harbored three proteomically distinct subpopulations, with differential protein expression analysis prioritizing differentially abundant NCL and MUCL1 as class markers. This proteomic fingerprint was subsequently validated by immunofluorescence on bone marrow sections, resulting in spatially distinct clusters linked to NCL or MUCL1 levels. We supplemented our molecular findings by qualitative scoring of patient samples by pathologists, which were in agreement with the observed morphological changes in the nuclear size between classes and match proteomically defined differences in nuclear makeup and stemness. Taken together, these findings establish SCP-MS as a powerful tool for uncovering functional heterogeneity in pediatric B-ALL with potential therapeutic and clinical relevance.

While flow-cytometry remains the clinical standard for immunophenotyping leukemic blasts, it carries some technical limitations in profiling molecular signatures between subpopulations within patients.

As demonstrated here, clinical flow-cytometry of all three patients had limited ability to reveal distinct subpopulations within the leukemic compartment (**Fig. 1B)**, consistent with known limitations of marker-based approaches in resolving deeper molecular heterogeneity^35^that can only partially be overcome by increasing the complexity of the flow-cytometry panel. Bulk proteomics on the same sorted samples was able to distinguish between patients but collapsed intra-patient variability into a single averaged signal (**Fig. 1C**), a well-recognized limitation of ensemble measurements in heterogeneous tumor samples^6^. By contrast, SCP-MS allows for resolving patient-specific subpopulations in an untargeted manner and, within B-ALL-3, identified three clusters with distinct proteomic identities that would have been entirely lost in a bulk experiment (**Fig. 2C**). This highlights a key advantage of single-cell resolution: the ability to detect rare or minority cell states that may be disproportionately relevant to disease outcomes^36,37^.

While single-cell transcriptomics has been applied to B-ALL to characterize leukemic cell states and their association with genetic alterations and clinical outcomes^38^, transcript abundance is an incomplete proxy for protein levels, with numerous studies demonstrating weak mRNA-protein correlation across cell types and conditions^17–19,39^.

Among the top differentially abundant proteins between subpopulations in B-ALL-3, Nucleolin (NCL) and Mucin-like protein 1 (MUCL1) stood out as candidate markers to differentiate them (**Fig. 3A**). NCL is a multifunctional nucleolar protein involved in ribosome biogenesis, cell proliferation, Wnt signaling activation, and anti-apoptotic signaling, and has previously been shown to be overexpressed in ALL where it directly promotes chemotherapy resistance and correlates with poor overall survival and relapse-free survival^40^. Furthermore, NCL has been identified as aberrantly active in hematopoietic stem- and progenitor cells, where it amplifies long-term culture-initiating cells and promotes execution of the stem-cell gene expression program through activation of Wnt signaling^41–43^. The co-occurrence of elevated NCL with high stemness scores^34^ in pClass 3 (**Fig. 3J**) is therefore consistent with a biologically aggressive, therapy-resistant phenotype. MUCL1, while less characterized in hematological malignancies, has been shown to promote tumor cell proliferation, colony formation, and epithelial-to-mesenchymal transition through Bcl-2 family upregulation and β-catenin deregulation, establishing it as an oncogenic driver in solid tumors^44,45^. Its emergence as a top differentially abundant protein in pClass 1 of B-ALL-3 warrants future investigation into its role in leukemic biology.

We were able to corroborate several facets of our SCP-MS findings using IHC and clinical pathology approaches. First, mapping of SCP-MS derived abundance classes onto segmented IHC data resulted in distinct spatial regions with differences in cell morphology. These NCL / MUCL1 abundance classes observed by SCP-MS did align with the measured nuclear size differences between them **(Fig. 3D-E; Supplementary Fig. 3D,E)**. Second, we observed differences in nuclear size when analyzing different blast morphologies annotated by clinical pathologists as well as in aspirate samples which served as the sample input for the SCP-MS for the first place **(Fig. 3G,H; Supplementary Fig. 3F,G)**.

While these results represent a promising proof-of-principle for extending clinical flow-cytometry analysis with SCP-MS, several limitations should be acknowledged. First, throughput remains a practical constraint of current SCP-MS workflows. First, the acquisition rate of approximately ∼40 cells per day limited the total number of cells analyzed to ∼410 across seven samples. While this is sufficient for proof-of-concept subpopulation discovery, it precludes robust statistical characterization of ultra-rare subpopulations and limits generalizability. Strategies to increase the acquisition rate or multiplex cells are actively developed^46,47^. While these will likely reduce this limitation in the future, they have so far only been demonstrated on larger cells. Second, protein depth is affected by the small size of lymphocytes, which contain less total protein than larger cell types. Our median of ∼1,000 proteins per cell represents approximately the top 10% of the expressed proteome, meaning that lower-abundance regulatory proteins, including many transcription factors and signaling intermediates, remain below the detection limit. Third, missing values remain an inherent feature of SCP-MS data due to stochastic sampling at the single-cell level. Although we applied kNN imputation to enable downstream analyses, imputed values introduce uncertainty and should be interpreted with caution, particularly for proteins with high missingness rates. Finally, our cohort currently consists of three B-ALL patients, limiting the ability to draw broad conclusions about subpopulation prevalence or clinical associations across the broader disease spectrum.

Future improvements in instrumentation will allow for faster and deeper molecular profiling of larger patient-cohorts in longitudinal studies^21^. For example, MRD and relapse-timepoint sampling of cohorts will allow tracking of subpopulation dynamics through disease evolution, an approach that has proven informative in bulk proteomics studies of pediatric ALL^10^. Additionally, integration of SCP-MS with single-cell RNA sequencing and spatial approaches will enable direct comparison of protein and transcript heterogeneity within the same cells and its environment, helping to illuminate mechanisms on several levels of regulation and co-dependencies. Finally, expanding the cohort to include patients stratified by molecular subtype, MRD status, and relapse outcome will be essential to evaluate whether the proteomic subpopulation signatures identified here have prognostic value.

Despite these stated limitations, this study demonstrates that SCP-MS is feasible directly on primary pediatric B-ALL material. Our workflow was able to identify functionally distinct proteomics subpopulations within individual patients that were invisible to both flow-cytometry and bulk proteomics. Moreover, these patient-specific subpopulations were orthogonally corroborated by spatial IHC and review of tissue- and aspirate-samples by a trained pathologist. Taken together, these results position SCP-MS as a valuable addition to the pediatric B-ALL research toolkit, ready to expand into larger, longitudinal patient cohorts to help define clinically relevant subpopulation at the molecular level.

## Acknowledgements

We gratefully acknowledge the work and staff of the BC Children’s Hospital BioBank for their assistance with sample and clinical data acquisition. We also would like to thank the patients and their families for generously participating in this study, without whom this research would not have been possible.

## Author contributions

Conceptualization: F.J. P.F.L.

Methodology: F.J., A.C., F.K., P. F. L.

Experimental Investigation: F.J., A.S.

Clinical & Pathology: A.S., J.W.B.

Computational Investigation: A.C., F.J. F.K.

Visualization: F.J., A.C., F.K.

Funding acquisition: P. F. L.

Supervision: F.K., A.S., P. F. L.

Writing – original draft: F.J., F.K., A.C., P. F. L.

Writing – review & editing: F.J., F.K., A.C., A.S., J.W.B., P. F. L.

## Materials and Methods

### Patient & Control Samples

#### Ethics statement

Patient specimens were collected by the Biobank staff at BC Children’s Hospital with informed consent obtained from patients and their parents during routine clinical care. All collections and experiments were conducted under approval from the University of British Columbia Children & Women’s Research Ethics Board (H17-01860) and in accordance with the ethical principles outlined in the WMA Declaration of Helsinki and the Belmont Report of the U.S. Department of Health and Human Services. All patients in the cohort were female, self-reported by patients or their parents.

#### Samples

Bone marrow samples from three different B-ALL patients were collected through the BCCHR biobank. We also obtained two healthy bone marrow aspirates and two peripheral blood samples from non-cancer pediatric patients.

### Flow-cytometry

Fresh bone marrow specimens obtained at diagnosis from subjects B-ALL-1,2,3 were analyzed using multicolor flow-cytometry according to the previously described clinical protocol at BC Children’s Hospital48. Specimens from subjects B-ALL-1 and B-ALL-2 were analyzed using a 10-color panel comprising CD58–FITC, CD49f–PE, CD20–ECD, CD38–PC5.5, CD19–PC7, CD33–APC, CD34–APC–Alexa Fluor 700, CD10–APC–Alexa Fluor 750, CD123eFluor 450, and CD45–Krome Orange. Because subject B-ALL-3 was analyzed at an earlier time point, its diagnostic specimen was assessed using the preceding 5-color clinical panel, comprising CD45–PC5, CD38–FITC, CD10–ECD, CD20–PE and CD19–PC7. Antibodies were used at dilutions established by prior titration and validated for clinical use.

| Antibody-fluorochrome combination | Vendor | Catalog # | Clone |
| --- | --- | --- | --- |
| CD58-FITC | Beckman Coulter | IM1218U | AICD58 |
| CD49f-PE | Invitrogen | 12-0495-81 | eBioGoH3 |
| CD20-ECD | Beckman Coulter | IM3607U | B9E9 |
| CD38-PC5.5 | Beckman Coulter | A70205 | LS198-4-3 |
| CD19-PC7 | Beckman Coulter | IM3628U | J3-119 |
| CD13/33-APC | Beckman Coulter | IM2471U | D3HL60.251 |
| CD34-APC AF700 | Beckman Coulter | A86354 | 581 |
| CD10-APC AF750 | Beckman Coulter | A89310 | ALB1 |
| CD123-eFluor450 | Invitrogen | 48-1239-42 | 6H6 |
| CD45-KRO | Beckman Coulter | A96416 | UCHT1 |

Following incubation for 15 min at room temperature in the dark, samples were processed using IntraPrep reagent (Beckman Coulter) according to the manufacturer’s instructions, washed, and resuspended for acquisition. Data were acquired on a Navios flow cytometer (Beckman Coulter), with 100,000 total events collected per tube. Flow-cytometry data analysis, including compensation adjustments, were done using Kaluza software (Beckman Coulter). Debris was excluded based on forward and side scatter characteristics, and doublets were excluded using forward-scatter area versus height. Leukemic blasts were identified as a CD45-dim, low-side-scatter population, with their identity confirmed by the expression pattern of the remaining markers in the panel.

### Mass Spectrometry sample preparation, cell sorting, and digestion

All five bone marrow samples (BMMCs) and two peripheral blood samples (PBMCs) were prepared for cell sorting by pelleting all the cells first (1200g for 5 min) followed by three times washing with Phosphate Buffered Saline (PBS). A portion of each sample was taken for negative control (unstained controls), and the remaining cells were incubated with 5 µL of the FITC anti-human CD19 Antibody [0.5 mg / mL] (Biolegend: 302205) in 95 µL of PBS. The staining was done in the dark at 4°C for 40 min and followed by three washes with PBS to ensure removal of the staining solution.

After cell counting (Froggabio, Denovix CellDrop), each sample was diluted to 10,000 cells / mL, ensuring optimal loading and sample processing. Prior to single cell dispensing 1.5 µL of lysis and digestion buffer [0.02% DDM (n-dodecyl-β-d-maltoside),1:30,000 of 10x iRT (Biognosys; Indexed Retention Time) and 1 ng / µL trypsin in 200 mM TEAB were deposited into a low binding 96 well plate using the automatic liquid dispenser iDOT (Dispendix). Using a Pala single cell dispenser (Bio-Techne), 1 cell / µL was dispensed into the prepared 96 well plates. The plates were centrifuged at 1,200g for 5 min and incubated at 37°C for 2 h in a humidified cell culture incubator.

### Liquid Chromatography and Mass Spectrometry (LC/MS)

Plates were dried using a SpeedVac (ThermoFisher; SPD120) and loaded in a randomized fashion into a Nanoelute 2 liquid chromatography system (Bruker Daltonics). Single cell peptides were resuspended in 2 µL of buffer A (0.1% formic acid (FA) in water) using the liquid handling functions of the Nanoelute 2.

The full 2 µL volume was loaded and the peptides separated on a 15 cm Elite Aurora reverse-phase C18 column with an inner diameter of 75 µm, particle size of 1.7 m, a pore size of 120 Å, and a 10 µm tip fused emitter (IonOpticks). Elution was performed with a flow rate of 300 nL / min and a gradient of 2 to 26% buffer B (0.1% FA in Acetonitrile (ACN)) over a 19 min gradient. This LC setup was coupled to a timsTOF Ultra Mass Spectrometer (Bruker Daltonics).

The Nanoelute 2 LC system was connected into a CaptiveSpray ESI source with a column-fused 10 µm emitter with 1500 V electrospray potential. Full MS data were acquired in the range of m/z 100-1700 and 0.6-1.45 1/K0 [V·s/cm2] in diaPASEF mode. A variable window width method was created using pydiAID31; 40 DIA windows over the range of 300–950 m/z were acquired with ramp and accumulation times of 100 ms (100% duty cycle) and 10 MS/MS ramps. Specific window placements are provided in **Supplementary Data 2**. The improved variable window (pydiAID31) method gave +40% protein IDs (**Supplementary Fig. 1B**). The estimated cycle time was 1.07 s. The collision energy gradually increased as a function of increasing mobility starting from 20 eV at 0.6 1/k_0_ to 59 eV at 1.6 1/k_0_.

### Preparation of Formalin-Fixed Paraffin-Embedded (FFPE) bone marrow sections for IHC

Bone marrow biopsies were obtained through the clinical lab. Biopsies were fixed in B-Plus fixative for 3 hours, then washed in water for 5 min. The biopsies were then decalcified in Rapidcal decalcification solution for 75 min, washed in water for 10 min, then placed in 10% neutral-buffered formalin. The biopsy was then processed overnight, embedded in paraffin wax, and sectioned at the microtome at 4 µm for IHC.

### Morphological classification and scoring of blasts in FFPE BM samples

The corresponding diagnostic bone marrow aspirate and hematoxylin-and-eosin-stained FFPE core biopsy were retrospectively reviewed by a hematopathologist to assess for morphologic heterogeneity. Blasts were classified according to cell size and nuclear chromatin characteristics, and the relative proportions of the morphologically distinct populations were estimated in representative, adequately preserved, blast-rich areas. The relative proportions of the two populations were estimated by evaluating representative, adequately preserved, blast-rich areas of the biopsy. Areas affected by crush artifact, necrosis, hemorrhage, or poor preservation were excluded.

### Immunofluorescence staining and mounting

FFPE tissue sections mounted on slides were baked at 65°C for 20 min, deparaffinized in xylene, and rehydrated through graded ethanol washes to distilled water. Antigen retrieval was performed in antigen retrieval buffer (Sodium Citrate, pH6) using steam heating in a rice cooker for 20 min, followed by cooling to room temperature. Sections were washed in PBS (3 x 5 min). Slides were then permeabilized with 0.2% Triton X-100 in PBS for 10 min and blocked in 2% normal goat serum (NGS), 0.02% Triton X-100, and 1% BSA in PBS for 1 h at room temperature. Directly conjugated primary antibodies were diluted in blocking buffer and incubated overnight at 4°C in a humidified chamber. The following antibody dilutions were used: anti-CD19 (1:50), anti-nucleolin (1:25), and anti-MUCL1 (1:25). After PBS washes, nuclei were counterstained with DAPI (1:50,000 in PBS) for 10 min at room temperature protected from light. Sections were washed with PBS and mounted using ProLong− Diamond Antifade Mountant (Invitrogen, cat. #P36961).

### Fluorescence Microscopy

Immunofluorescently labelled cells were imaged at room temperature using a ZEISS Celldiscoverer 7 automated widefield fluorescence microscope equipped with a Plan-Apochromat 5x/0.35, Plan-Apochromat 20x/0.7 Autocorr, Plan-Apochromat 20x/0.95 Autocorr and Plan-Apochromat 50x/1.2 W Autocorr objective lenses (ZEISS). Fluorescence was excited sequentially using LED illumination (385, 470, 567 and 625 nm). The system automatically detected vessel bottom material and thickness and applied spherical aberration correction via the motorized Autocorr mechanism. Images were acquired with a ZEISS Axiocam 807 mono camera using ZEN blue software (ZEISS).

Bone marrow samples were imaged using the Plan-Apochromat 20x/0.7 Autocorr objective at a final pixel size of 0.225 µm (4.5 µm sensor pixel). Three fluorescence channels were acquired sequentially per field: DAPI (ex 353 nm / em 465 nm; 40 ms, 30% LED), R-PE (NCL; ex 565 nmm/ em 576 nm; 300 ms, 50% LED) and Alexa Fluor 647 (CD19; ex 653 nm / em 668 nm; 200 ms, 30% LED). Samples were imaged as tiled acquisitions (21 x 35 tiles, 10% overlap, meander scan) covering approximately 13.7 x 15.5 mm per sample, with automated focus surface interpolation. All tiles were acquired as single focal planes. Identical acquisition settings were applied across all samples.

### Image analysis

Image analysis was performed in Python (segmentation, registration) and R (statistics, visualization). CZI pyramid images were exported per channel as 8-bit greyscale .tiff files. Analysis comprised: (1) segmentation, (2) QC filtering, (3) spatial analysis, (4) NCL-MUCL1 co-registration and spatial class analysis (pt3), (5) H&E integration, (6) statistics and visualization.

#### (1) Segmentation

Nuclei were segmented with CellposeSAM (cpsam_20260504_BM) and NCL signal within nuclear masks measured (mean, integrated intensity). Tiles with <5% non-zero pixels were excluded as empty. Nuclear masks were expanded by distance transform, capped at the midpoint between neighboring nuclei to avoid double-counting. CD19 status per nucleus was called from tile-mean (I_avg) and nuclear (I_nuc) CD19 intensity: positive if I_nuc > I_avg ™ 0.5*SD(I_avg). QC overlays (segmentation, CD19 calls, NCL signal) were saved per tile.

#### (2) QC filtering

Tiles were filtered on nuclear count via 3*MAD, retaining median ± 3*MAD, with a hard floor of 250 nuclei/tile to exclude near-empty tiles.

#### (3) Spatial analysis

Nuclei were assigned global canvas coordinates by stitching tiles using embedded Zeiss coordinate strings. NCL intensity was tile-corrected for acquisition offset (ncl_corrected = ncl_mean ™ tile_mean + global_mean), and CD19 status re-called globally on corrected intensity. Nuclei were binned into eight percentile classes (P0-5 to P95-100) and a four-class scheme (P0-50, P50-75, P75-90, top 10%) by corrected NCL rank across all tiles.

#### (4) NCL-MUCL1 co-registration and spatial class analysis

NCL and MUCL1 sections (consecutive biopsies) were registered without fiducials, using nuclear outlines from 10x-downsampled stitch canvases (2.25 µm/pixel), in three stages: (i) coarse translation by centroid shift, refined via phase cross-correlation (ii) ICP rigid registration on nucleus centroids, weighted toward large nuclei to limit z-section sensitivity, with per-iteration outlier rejection; (iii) TPS deformable warp using ICP-matched centroids as control points. The transform was applied to MUCL1 coordinates; nuclei outside the NCL canvas were flagged.

SCP-informed classes were defined on co-registered IHC using thresholds: Class 1 (MUCL1-high/NCL-low), Class 2 (neither high), Class 3 (NCL-high/MUCL1-low); double-positive cells were retained but excluded from class comparisons. Class and nuclear area were compared pairwise by Wilcoxon rank-sum with Benjamini-Hochberg correction.

To reduce sensitivity to registration error (∼5 µm) and slice Δz, cells were also aggregated into 10×10 µm spatial blocks, with per-block mean NCL, MUCL1, DAPI and CD19 positivity computed and classified using the same thresholds.

#### (5) H&E integration

A whole-slide H&E image was registered to the NCL coordinate space for morphological context. Hematoxylin centroids were registered to NCL/DAPI centroids using the same ICP+TPS pipeline, and the transform applied to single-cell and block coordinates. Per-tile overlays rendered SCP classes on H&E; annotations were exported as GeoJSON (one file per class) for QuPath review.

#### (6) Statistics and visualization

Analyses and figures were generated in R. Group comparisons used Wilcoxon rank-sum tests, pairwise SCP-class comparisons used Benjamini-Hochberg-corrected Wilcoxon tests, and effect sizes are reported as group medians. Figures were exported as Cairo PDF with source data CSVs alongside.

### Proteomics data analysis

Proteomic raw data was searched in directDIA mode using Spectronaut Pulsar X (Biognosys V19.3)49 using a human SPROT reference proteome without isoforms (20,656 sequences) FASTA downloaded from UniProt on July 17th, 2024. For the search, enzyme, and digestion type were set to Specific, and Trypsin/P, acetyl (Protein N-term), and oxidation (M) were set as variable modifications with no fixed modification. Maximum and minimum peptide length were set to 7 and 52 amino acids, respectively, and missed cleavage was set to 2. Precursor and protein FDR were set to 1%, and a minimum of 2 peptides were used for quantification. Protein group reports were exported from Spectronaut and included the variables PG.ProteinGroups, PG.Genes, PG.ProteinNames, PG.Descriptions, and PG.Quantity, which was used as the quantitative protein abundance measure. Data analysis and visualization were performed using R v4.5.1 within RStudio v2026.01.0+392, using the tidyverse ecosystem (ggplot2, dplyr, tidyr).

#### Bulk Proteomics

Three technical replicates from each patient were analyzed. No samples were excluded during quality control; therefore, all three replicates from each patient were retained for downstream analyses. Across the dataset, a median of 3,831 proteins was identified per sample. Where required, missing quantitative values were imputed using a k-nearest neighbours (kNN) algorithm (k = 5), applied separately within each patient. Hierarchical clustering was performed using Euclidean distance in the ComplexHeatmap package in R. Reproducibility between technical replicates was further assessed by pairwise Spearman correlation analysis using the non-imputed quantitative protein abundances.

#### Single Cell Proteomics

Single cells with a low number of identified proteins (<400) were considered low quality and therefore excluded from downstream analysis. After filtering, a total of 385 cells were retained, comprising Patient 1 (n = 84), Patient 2 (n = 80), Patient 3 (n = 70), BMMC.1 (n = 46), BMMC.2 (n = 45), PBMC.1 (n = 30), and PBMC.2 (n = 30). Across the filtered dataset, a median of 928 proteins was identified across cells, corresponding to 1,412 unique proteins detected overall. Missing quantitative values were imputed using a k-nearest neighbours (kNN) algorithm (k = 5), implemented separately within each biological condition. The imputed dataset was used exclusively for unsupervised analyses requiring complete data matrices, whereas all statistical analyses were performed using the original non-imputed quantitative data. To investigate global similarities among single cells, hierarchical clustering was performed using Euclidean distance in the ComplexHeatmap package in R. Dimensionality reduction was carried out using Uniform Manifold Approximation and Projection (UMAP). Cellular trajectories were inferred using Partition-based Graph Abstraction (PAGA) tree analysis, where pClasseswere identified using k-means clustering.

Differential protein abundance analyses were conducted using linear models implemented in the limma package on the non-imputed data. Proteins were considered significantly different using cutoff parameters of |log_2_FC| > 1 and q-value < 0.05. Multiple testing corrections were performed using the Benjamini–Hochberg false discovery rate (FDR) method. Functional interpretation of differentially abundant proteins was performed by Gene Ontology (GO) over-representation analysis using the compareCluster function implemented in the clusterProfiler R package.

The Cancer-Associated Protein list was adapted from the pediatric oncology panel described by Lorentzian et al. (2023)10 and expanded to include additional therapeutically relevant proteins, including drug targets. The leukemia-related protein list was compiled based on multiple studies that performed ALL-specific analyses50–52. Both lists were used in selected downstream analyses.

### Data, Materials and Code availability

