## Supplemental Figures 1-3 for "Single-Cell proteomics discerns patient-specific subpopulations in pediatric B-cell acute lymphoblastic leukemia"

bioRχiv

Jayousi et al., 2026

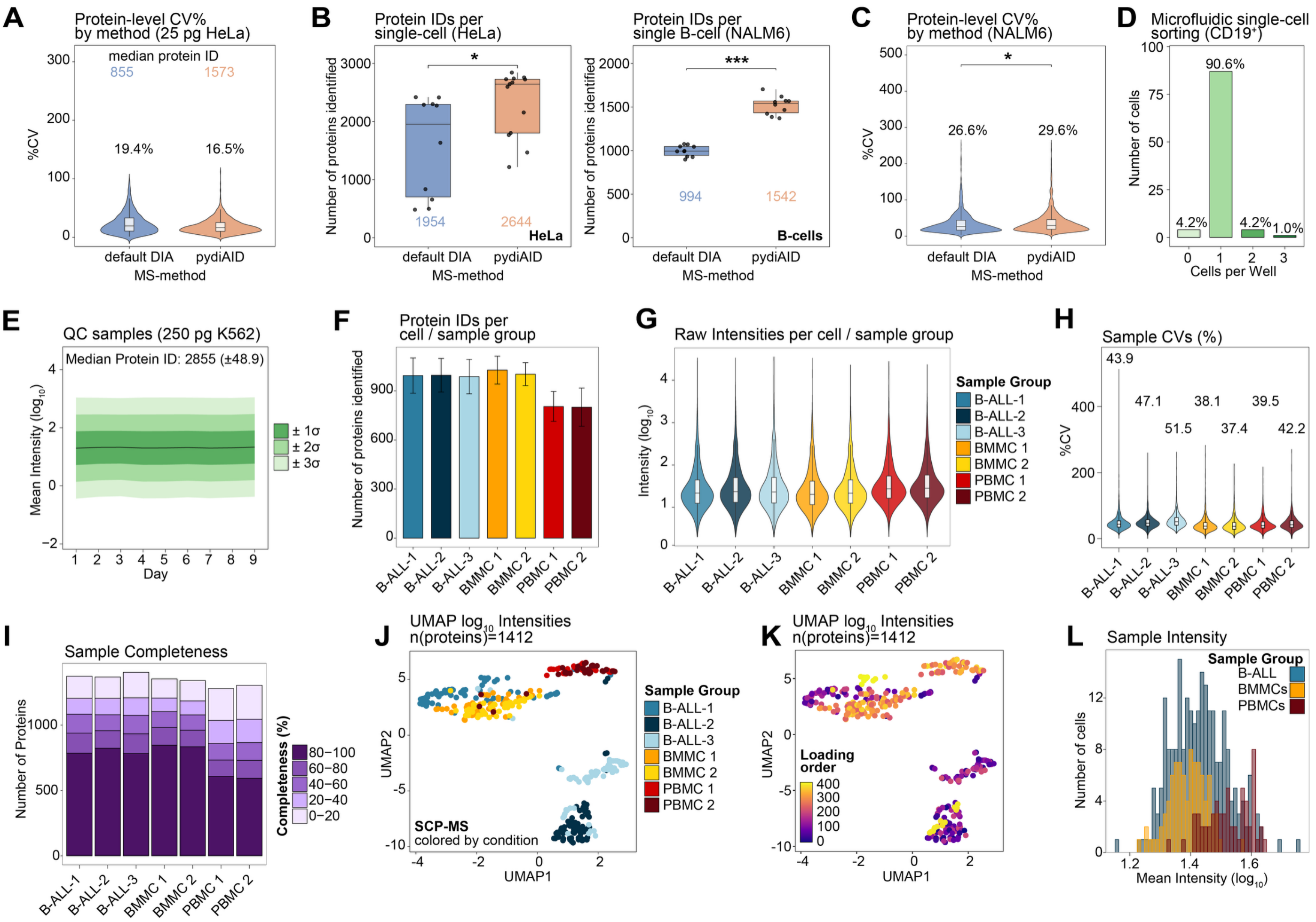


**Supplementary Figure 1. Quality control and data characteristics of single-cell proteomic measurements. (A)** Violin plot of cell-equivalent coefficients of variation (%CV) of HeLa dilutions using default (19.4% median %CV, 855 IDs) or pydiAID-optimized (16.5% median %CV, 1573 IDs) MS acquisition. **(B)** Boxplot of identified protein IDs per single-cell HeLa cell (left) or NALM6 cells (right) using default (blue, median = 1954 / 994 IDs [HeLa / NALM6]) and pydiAID-optimized MS acquisition (orange, median = 2644 / 1542 IDs [HeLa / NALM6]). 10 replicate single-cells were used per method and cell line. **(C)** Violin plot of protein-level %CV of single-cell NALM6 SCP-MS using default (26.6%) or pydiAID-optimized (29.6%) MS acquisition. 10 replicate single-cells were used per method and cell line. **(D)** Barplot showing isolation accuracy of microfluidic single-cell sorter per well. **(E)** $Log_{10}$ Signal intensity of K562 QC standards over 9-day SCP-MS acquisition window. **(F)** Barplot of number of identified proteins per cell across patient and control samples. Mean and standard deviation (SD) are shown for each condition. **(G)** Violin plot of raw intensities ($log_{10}$) across sample groups. **(H)** Distribution of protein %CV across cells for each condition. **(I)** Stacked bar graph depicting protein completeness across cells, shown as the proportion of quantified proteins within completeness intervals. **(J)** UMAP embedding of single cells based on log10-transformed protein intensities across B-ALL patients, bone marrow mononuclear cell controls (BMMC), and peripheral blood mononuclear cell controls (PBMC). **(K)** As I, but loading orders of MS acquisition is mapped onto UMAP embedding. **(L)** Distribution of mean raw protein intensities ($log_{10}$) across conditions.

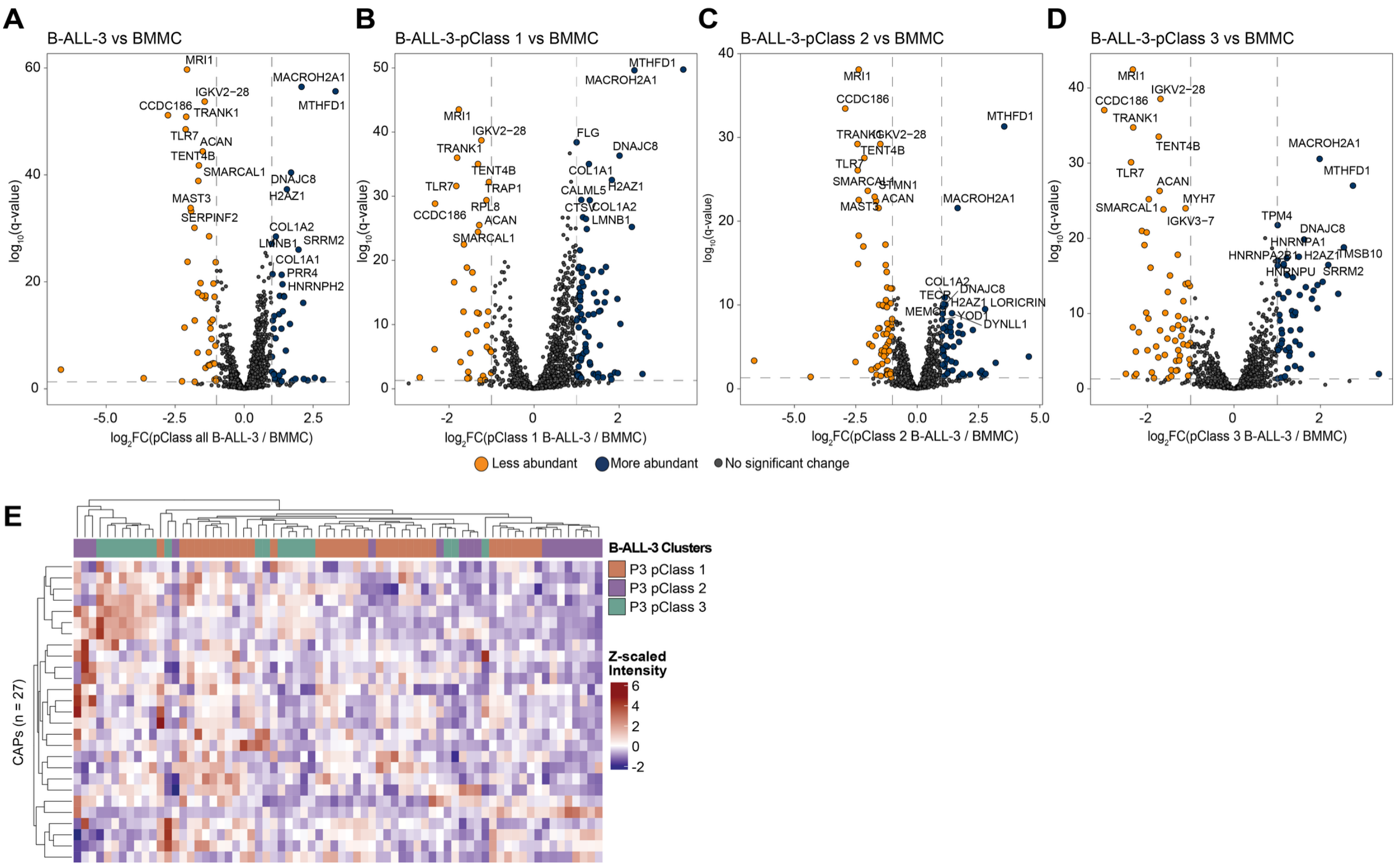


**Supplementary Figure 2. Analysis of B-ALL-3 clusters. (A-D)** Volcano plots of B-ALL-3 (A) all pClasses, (B) pClass 1 (B), (C) pClass 2 and (D) pClass 3 vs BMMCs. Top 10 hits are labeled. **(E)** Heatmap of Cancer-associated proteins (CAP)-abundance across B-ALL-3 pClasses 1-3, shown as z-scaled protein intensities.

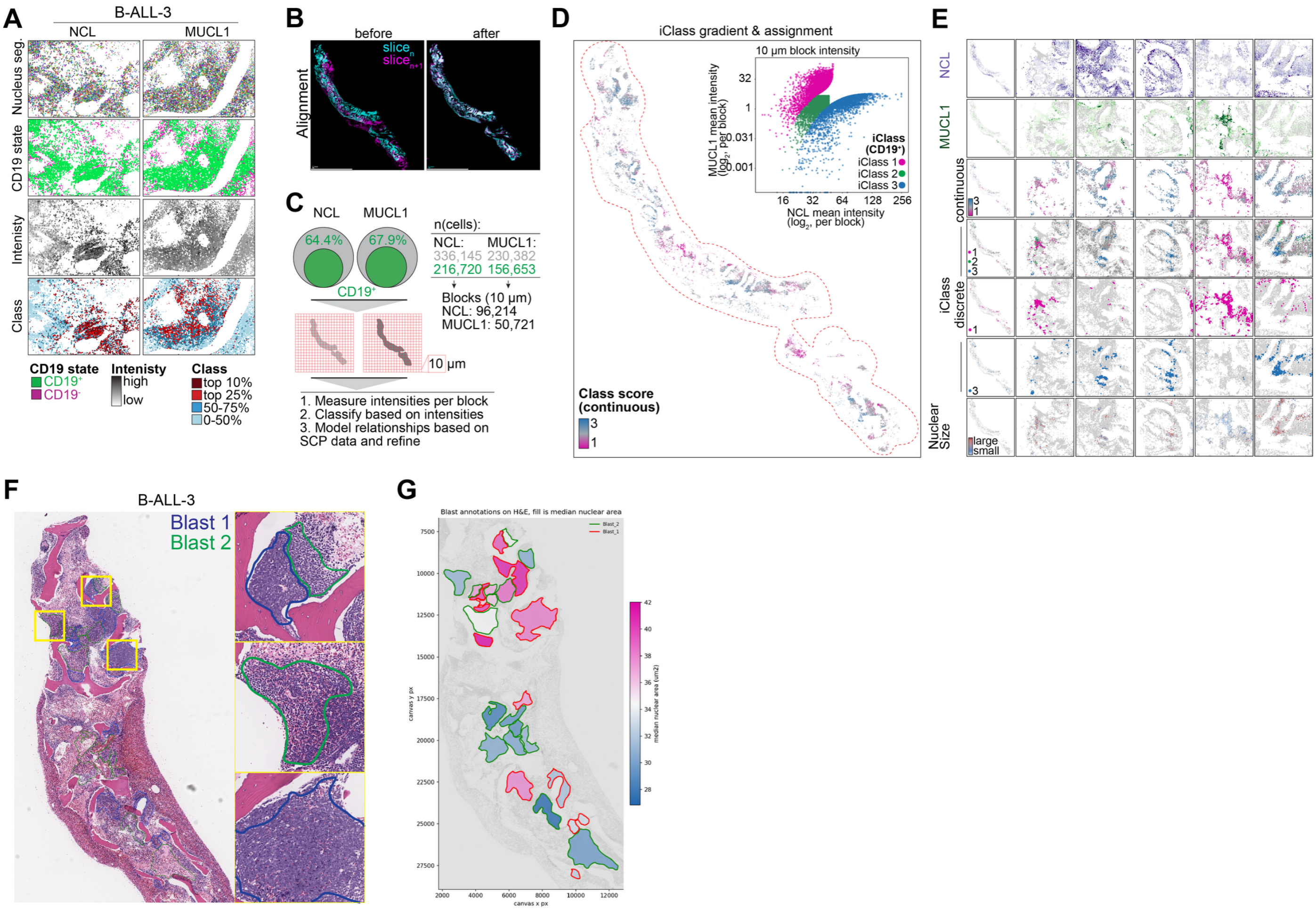


**Supplementary Figure 3: Strategy for SCP-MS result validation using immunohistochemistry, ML-aided image processing and pathology-scoring. (A)** From top to bottom: Example tiles from B-ALL-3 depicting the results of nuclear segmentation, CD19 classification, intensity per nuclear object and global classification for NCL (left) and MUCl1 (right). **(B)** Nuclear objects of consecutive slices shown before (left) and after global alignment (right). **(C)** Approach for the evaluation of IHC data from consecutive slices on 10 $\mu$m blocks-basis for integrating SCP abundance trajectories. **(D)** Spatial map of iClass score (continuous scale) applied on 10 $\mu$m blocks scale. Inset plots intensity of NUC x-axis vs MUCL1 (y-axis) per 10 $\mu$m block and are color coded by assigned iClass. **(E)** Example images depicting spatial heterogeneity of BM sample (from top to bottom): NCL intensity, MUCL1 intensity, iClasses continuous scale, overlay iClass 1-3, iClass 1, iClass 3, average nuclear size per block. **(F)** Overlay of H&E staining of B-ALL-3 with manual blast annotations made by pathologists. **(G)** Overlay of blast annotations colored according to mean nuclear size.
